# Enhancing predictive accuracy of yield traits in cassava through multi-trait genomic prediction

**DOI:** 10.64898/2026.07.01.735838

**Authors:** Gabriel Mamedio de Freitas, Diana C. Solarte Certuche, Jean-Luc Jannink, Eder Jorge de Oliveira, Antonio Augusto Franco Garcia

## Abstract

Multi-trait genomic prediction offers a practical route to improve selection for costly, complex traits in clonally propagated crops such as cassava. In a Brazilian breeding panel of 1,078 cassava clones genotyped with 25,923 SNPs and phenotyped for six agronomic traits, we compared single-trait (ST) and multi-trait (MT) GBLUP models. Stage-wise mixed models produced BLUEs that fed into ST and MT-GBLUP. We tested five cross-validation schemes that mimic breeder realities: ST baseline (CV1); naive all-traits MT prediction for unphenotyped candidates (CV2); MT prediction using auxiliary trait phenotypes in the test set (CV3); and two sparse-phenotyping regimes with missingness by trait (CV4) or by clone (CV5) at 25%, 50%, and 75% levels. The main results were that, under the ST baseline (CV1), predictive ability ranged from 0.50 for DMC and 0.45 for FRY down to 0.13 for Le.Dis. A naive full MT model (CV2) performed approximately on par with ST-GBLUP. In contrast, MT designs (CV3) that included informative auxiliary traits, such as shoot yield and combinations with plant vigor and leaf disease severity, yielded small gains for DMC with predictive ability of approximately 0.51 (+2%), while FRY predictive ability increased to approximately 0.65 (+44%), accompanied by RMSE reductions for FRY up to approximately 13.5% (e.g. RMSE approximately 6.2). Sparse-phenotyping simulations (CV4/CV5) demonstrated that MT models sustain or even improve predictive ability under realistic missing-data regimes ≈ 0.62λ0.65). Selection concordance between MT and ST top-10% sets was generally high (>0.80), and MT configurations produced measurable improvements in expected selection response and genetic gain per cycle for several target traits. These results indicate that strategically implemented MT-GBLUP, using a small set of biologically and operationally informative auxiliary traits and optimized sparse phenotyping, can materially increase predictive accuracy and selection efficiency for economically critical cassava traits while reducing phenotyping burden.

## Introduction

Cassava (*Manihot esculenta* Crantz), a member of the Euphorbiaceae family domesticated in South America (Ceballos, Iglesias, et al., 2004), is a cornerstone crop for food security across the tropics. Its resilience in marginal environments and its multiple end uses—fresh consumption, animal feed, and industrial processing make cassava a central element of many agri-food systems. Today, cassava is one of the world’s major sources of carbohydrates and underpins the livelihoods of millions across Africa and Asia (Ceballos, Rojanaridpiched, et al., 2020). In 2023, global production reached approximately 333.68 million tons, with Brazil producing about 18.51 million tons (FAOSTAT, 2025). In Brazil, cassava is cultivated nationwide and plays a dual role in subsistence and agro-industrial value chains, which motivates breeding priorities focused on root quality and industrial yield (Andrade, Sousa, Oliveira, et al., 2019; Oliveira, Resende, et al., 2012).

Conventional cassava breeding relies on recurrent phenotypic selection, crosses among elite clones, and multi-environment regional trials, but it faces intrinsic biological and operational constraints. Vegetative propagation, environmentally dependent flowering, long selection cycles, and the generally low heritability of several key yield traits make the pipeline slow and expensive, often spanning 8–10 years from cross to variety release (Ceballos, Iglesias, et al., 2004; Ceballos, Kulakow, and Hershey, 2012; Andrade, Sousa, Oliveira, et al., 2019). These constraints are particularly problematic for characteristics that are time-consuming or costly to measure, reducing the effectiveness of selection in the early stages.

Advances in molecular markers since the 1980s and the recent surge in genotyping capacity have transformed breeding strategies (Crossa et al., 2017; Ramesh et al., 2020). Although marker-assisted selection (MAS) provided useful early tools, its limited power in cassava, partly due to early QTL studies structured around highly contrasting crosses, motivated the adoption of genomic selection (GS), which uses genome-wide markers to predict breeding values without requiring prior QTL mapping (Meuwissen, Hayes, and Goddard, 2001; Nadeem et al., 2018; Wang et al., 2018). In cassava, GS enables early identification of superior clones, even in seedling stages, improves the prediction of complex or costly-to-measure traits, and offers a practical route to shorten breeding cycles and reduce phenotyping demands (Ceballos, Kulakow, and Hershey, 2012; Ceballos, Rojanaridpiched, et al., 2020; Oliveira, Resende, et al., 2012).

Single-trait GS (ST-GS) models have already demonstrated their efficacy in cassava to improve key yield components and resistance to major diseases like Cassava Mosaic Disease (CMD) and Cassava Brown Streak Disease (CBSD) (Wolfe, Rabbi, et al., 2016; Wolfe, Del Carpio, et al., 2017; Kayondo et al., 2018; Torres et al., 2019; Ozimati et al., 2018). Despite these advances, ST-GS models do not fully capture the complexity of breeding programs, which rarely focus on a single trait. Breeders typically select a combination of attributes simultaneously, often using selection indices. Multi-trait GS (MT-GS) aligns more closely with this reality by leveraging the genetic and residual covariances between traits (Okeke et al., 2017). This approach allows information from a highly heritable and correlated secondary trait to improve the predictive accuracy for a primary trait with low heritability, a principle well established in crops such as maize, wheat, and oat (Calus and Veerkamp, 2011; Jia and Jannink, 2012; Lado et al., 2018; Oliveira, Resende Jr, et al., 2020; Gill, Halder, et al., 2021; Gill, Brar, et al., 2023; Dhakal et al., 2024). Furthermore, MT-GS provides a powerful framework for predicting expensive or difficult-to-measure traits using data from easily scored auxiliary traits, particularly in scenarios with incomplete or imbalanced phenotyping (Lado et al., 2018; Matias et al., 2019).

This capability is especially relevant for cassava, where agronomic traits like plant vigor are easily evaluated, whereas key yield traits such as fresh root yield and dry matter content demand significant time, labor, and resources. However, the application of MT-GS in cassava breeding remains nascent. While a foundational study by Okeke et al. (2017) demonstrated the potential of multivariate models, it did not explore their utility in realistic breeding scenarios, such as leveraging secondary traits or addressing systematic missing data, where not all traits are phenotyped on all individuals. This represents a critical knowledge gap, as exploiting these scenarios could unlock significant efficiency gains in operational cassava breeding programs. Traits like fresh root yield and dry matter content, which are pivotal for both industrial and fresh consumption markets, often exhibit low-to-moderate correlations and are ideal candidates for enhancement through MT-GS frameworks (Joaqui Barandica et al., 2016).

Given this context, we hypothesized that MT-GS models would significantly outperform conventional ST-GS models in practical breeding scenarios characterized by partial phenotyping and the availability of correlated secondary traits. To test this hypothesis, this study was designed to compare the performance of ST and MT genomic prediction models for key traits in cassava across four realistic scenarios: (i) simultaneous prediction of six agronomic and yield traits for selection candidates with no phenotypic data; (ii) joint prediction of fresh root yield and dry matter content using correlated, easily measured secondary traits; (iii) prediction of fresh root yield and dry matter content when phenotypic data for the secondary traits are available but data for the primary traits are missing in the test population; and (iv) prediction of fresh root yield and dry matter content when entire clones have missing phenotypes for the primary traits, simulating the early-stage selection of genotyped but unphenotyped individuals.

## Methods & Materials

### Plant material, field trials and experimental management

We evaluated 1,078 cassava clones across eight field trials conducted at the Cassava Breeding Program of Embrapa Mandioca e Fruticultura located in Cruz das Almas (Latitude:12°48’38” S Longitude: 39°06’26” W) and Laje (Latitude: 1306’38.4” S Longitude:3916’20.4” W), Bahia, Brazil, between 2014 and 2020. The panel comprised improved clones from recurrent selection, plus varieties from research stations and local farmers. Planting took place between May and July during the rainy season. Plots were established at 0.90 m inter-row × 0.80 m intra-row spacing and standard agronomic practices for cassava cultivation were applied. All plants in each plot were harvested 11–12 months after planting. The following traits were recorded:

1. Dry Matter Content (DMC, %): estimated using the gravimetric method described by Kawano, Fukuda, and Cenpukdee (1987). Approximately 5 kg of clean roots, with excess soil removed and root ends trimmed, were weighed in air and subsequently in water. Dry matter content was then calculated as: 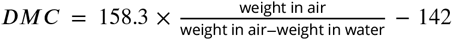 where weight in air and weight in water correspond to the root weights measured in air and water, respectively;
2. Fresh Root Yield (FRY, *t* ⋅ *ha*^−1^): calculated as the total fresh weight of harvested roots per plot. Only marketable roots, defined as roots free from significant pest and disease damage and conforming to commercial standards of size and shape, were included in the assessment;
3. Shoot yield (SHY, *t*.*ha*^−1^): total above ground biomass (leaves, petioles, stems);
4. Plant Architecture (Pl.Arch): scored 1–5 (scored on a 1 to 5 scale:5 = excellent (no branching or erect stems); 4 = good (branching above1.60 m or lower branches with at least 1.60 m of erect stem); 3 = moderate (branching above 1.20 m or lower branches with at least 1.20 m of erect stem); 2 = poor (branching above 0.80 m or lower branches with at least 0.80 m of erect stem); 1 = very poor (highly branched clones with less than 0.80 m of erect stem);
5. Vigor (Vigor): visual score at 1.5 months, 1 (low) to 5 (high);
6. Leaf Disease Severity (Le.Dis): visual 0–5 scale for *Passalora vicosae, Cladosporium henningsii* and *Passalora manihotis* (0 = clean plant, no infection; 1 = few affected leaves at the bottom; 2 = 50% of bottom leaves affected; 3 = symptoms on bottom and middle leaves with yellowing/defoliation below; 4 = high disease incidence throughout the plant; 5 = completely defoliated plant).

### Phenotypic analysis

Because trials were unbalanced across years and locations, we used a two-stage (stage-wise) analysis to: (*i*) adjust plot-level data for within-trial design and spatial effects, and (*ii*) combine trial-level estimates into across-environment estimates that account for differing precision among trial estimates. The two-stage approach yields environment-specific adjusted means (BLUEs) in stage 1 and then models these BLUEs in stage 2, weighting them by their precision. This procedure is standard in multi-environment trial (MET) analysis and supports downstream genomic analyses when full single-stage fitting is impractical (Smith, Cullis, and Gilmour, 2001).

Stage 1 (within-trial adjustments and spatial modeling): Within each trial, we fitted a linear mixed model to remove systematic, non-genetic effects and to obtain adjusted genotype means (BLUEs). The model was:

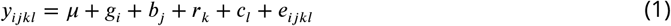

where *y*_*ijkl*_ is the plot observation for genotype *i* at the block *j*, row *k*, and column *l, µ* is the overall mean, *g*_*i*_ is the genotype effect (treated as fixed to obtain BLUEs), *b*_*j*_ is the random effect of the *jth* block with 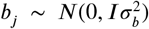, *r*_*k*_ and *c*_*l*_ are random row and column or spatial terms (the latter modeled using penalized tensor-product marginal B-splines where appropriate), and *e*_*ijkl*_ is the residual error with 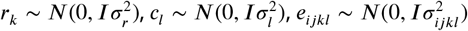 where *σ*_*r*_, *σ*_*c*_ and *σ*_*e*_ represents row, column and residual variances, respectively. Spatial correction using tensor-product B-splines was performed according to Rodriguez-Alvarez et al. (2018). This stage produced environment-specific genotype BLUEs and their associated standard errors for each trait and trial.

Stage 2 (cross-environment models and precision-weighted analysis): The BLUEs obtained in Step 1 were combined across trials using mixed models that explicitly accounted for the effects of genotype, environment, year, and their interactions. In this study, the environment was defined as the combination of location (experimental site) and year. Two model structures were applied according to the trait class:

Model 1 (Plant vigor and architecture — mainly categorical/score traits):

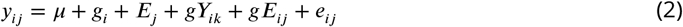

Model 2 (Dry matter content, fresh root yield and shoot yield — continuous, yield/quality traits):

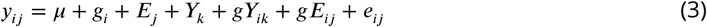

where *y*_*ij*_ phenotypic value of the *i*_*th*_ genotype in the *j*_*th*_ environment, *µ* is the overall mean, *g*_*i*_ is the fixed effect of the *i*_*th*_ clone, *E*_*j*_ is the random effect of *j*_*th*_ environment, *Y*_*k*_ is the random effect of the *k*_*th*_ year, *gY*_*ik*_ is the random effect of the interaction between the *i*_*t*_*h* clone and the *k*_*th*_ year, *gE*_*ij*_ represents the random effect of the interaction between the *i*_*th*_ clone and the *j*_*ij*_ environment, and *e*_*ijk*_ ~ *N*(0, *W*) with *W* is a diagonal matrix containing the inverse of the standard errors of the BLUEs from the first stage 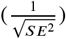. These standard errors were used to scale the residual variance in the second stage analysis (Smith, Cullis, and Gilmour, 2001; Frensham, Cullis, and Verbyla, 1997). This variance-weighted stage-2 fitting accounts for heterogeneity of BLUE precision among trials and is recommended practice for stage-wise MET analysis. The random effects follow the distributions: 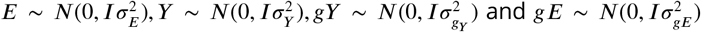 where 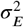 represents environmental variance, 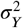 represents year variance, 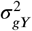 represents the variance due to the interaction between genotype and year, and 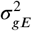 represents the variance due to the interaction between genotype and environment.

For leaf disease severity, we fitted a trial-level mixed model that preserves plot-level structure and spatial terms:

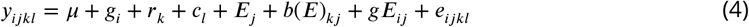

where *y*_*ijkl*_ is the phenotypic observation of the *i*_*th*_ genotype evaluated in the *j*_*th*_ environment, within the *k*_*th*_ row and *l*_*th*_ column; *µ* is the overall mean; *g*_*i*_ is the fixed effect of the *i*_*th*_ genotype; *r*_*k*_ is the random effect of the *k*_*th*_ row; *c*_*l*_ is the random effect of the *l*_*th*_ column; *E*_*j*_ is the random effect of the *j*_*th*_ environment; *b*(*E*)_*kj*_ is the random effect of the *k*_*th*_ block within the *j*_*th*_ environment; *gE*_*ij*_ is the random genotype-by-environment interaction effect; and *e*_*ijkl*_ is the residual error term. The random effects were assumed to follow independent normal distributions: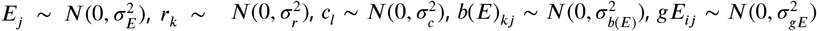, and 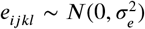, where 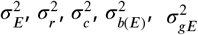, and 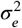 denote the variance components associated with environment, row, column, block-within-environment, genotype-by-environment interaction, and residual error, respectively.

### Variance components, heritability and testing of random effects

After obtaining stage-two BLUEs, genotypes were modeled as random effects to estimate genetic variance components for downstream calculations and genomic analyses. Random effects were evaluated by likelihood ratio tests (LRT): nested full and reduced models were compared and LRT statistics assessed with *a*^2^ distribution and appropriate degrees of freedom; effects were considered significant at *α* = 0.05 unless otherwise stated.

We estimated broad-sense heritability (*H*^2^) on an entry-mean basis following (Cullis, Smith, and Coombes, 2006), using the mean variance of pairwise differences of genotype BLUPs:

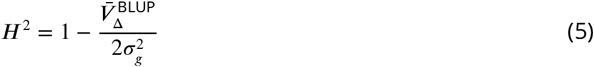

where 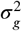 the estimated genotypic variance and 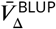 is the mean variance of the difference between two BLUPs (Cullis, Smith, and Coombes, 2006) and the narrow-sense heritability (*h*^2^) was estimated following

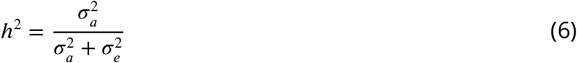

where 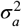 and 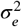 represent the additive and residual components based on markers.

Variance components and broad (*H*^2^) and narrow-sense (*h*^2^) heritability estimates were derived from mixed models fitted in `ASReml-R` (Butler et al., 2009) and `BGLR` (Campos et al., 2017), respectively. Pairwise Pearson correlations among traits were computed using the stage-two BLUEs.

### Genotypic data

DNA extraction and quality control were performed by Intertek (Australia). Genomic libraries were constructed using the DArTseq complexity reduction protocol (Kilian et al., 2012) at Diversity Arrays Technology (Canberra, Australia). Sequencing was carried out on an Illumina HiSeq 2500 platform (Illumina, USA). Raw reads underwent barcode removal and trimming using the `STACKS` pipeline (Catchen et al., 2013). Trimmed reads were aligned to the Cassava Reference Genome *v*6.1 using `BWA` (Li, 2010). Variant calling was conducted using GATK (Auwera and O’Connor, 2020), excluding indels and multiallelic sites. Quality filtering followed the `TASSEL` GBS 5.0 pipeline v 5.2.90 (Glaubitz et al., 2014), with thresholds: minimum call rate = 80 %, minor allele frequency (MAF) 1 %. Missing genotypes were imputed using the LD-KNNi approach (Money et al., 2015). The resulting dataset comprised 25,923 informative SNPs for downstream genomic prediction.

### Single-trait and multi-trait genomic prediction models

For each trait, we fitted a univariate Genomic Best Linear Unbiased Prediction (GBLUP) model:

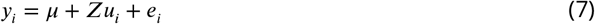

where *y*_*i*_ is the vector of BLUEs for trait *i, µ* is the overall intercept, and *Z* is the incidence matrix relating observations to genomic effects. The vector of additive genomic effects was assumed to follow 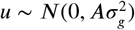, where *A* is the genomic relationship matrix and 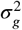 is the additive genomic variance. The residual errors were assumed to follow 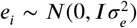, where 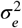 represents the residual variance.

We extended the GBLUP model to a multivariate context, enabling joint modeling of multiple traits with genetic covariances. The full model can be expressed in matrix form as:

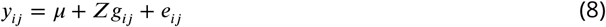

where 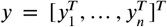 is the vector obtained by concatenating the BLUE vectors for all *n* traits; 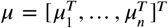 is the vector of overall means for each trait; *Z*_*n*_ is a block-diagonal incidence matrix relating genotypes to trait-specific genomic effects; 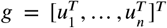 is the vector of stacked additive genomic effects across traits; and 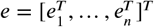 is the vector of stacked residual effects.

The additive genomic effects were assumed to follow *g ~ N*(0, *G* ⊗ *A*), where *A* is the genomic relationship matrix and *G* is an unstructured *n* × *n* genetic variance-covariance matrix, with diagonal elements representing genetic variances and off-diagonal elements representing genetic covariances among traits. Residual effects were assumed to follow *e ~ N*(0, *I* ⊗ *R*), where *R* is a diagonal residual variance-covariance matrix, assuming independence of residuals among traits.

Both ST-GBLUP and MT-GBLUP models were implemented using the `BGLR` and `Multitrait` func-tions from the `BGLR` package. Posterior distributions of genomic effects and variance components were estimated using 30,000 total MCMC iterations with 5,000 for burn-in. The genomic relationship matrix (*A*) was computed according to VanRaden (2008) using the `Gmatrix` function implemented in the `AGHmatrix` package (Amadeu et al., 2023).

### Cross-validation schemes

To assess consistency and predictive performance, we defined multiple cross-validations (CV) scenarios, illustrated in Figure 1. Each cross-validation (CV) sampling scheme was repeated 100 times. In each replicate, clones were randomly partitioned into: i) a training set comprising 80% of the clones, for which both genotypic and phenotypic data were used to fit the genomic prediction models; and ii) a test set comprising the remaining 20% of the clones, for which only genotypic data were used to estimate GEBVs.

**Figure 1.**
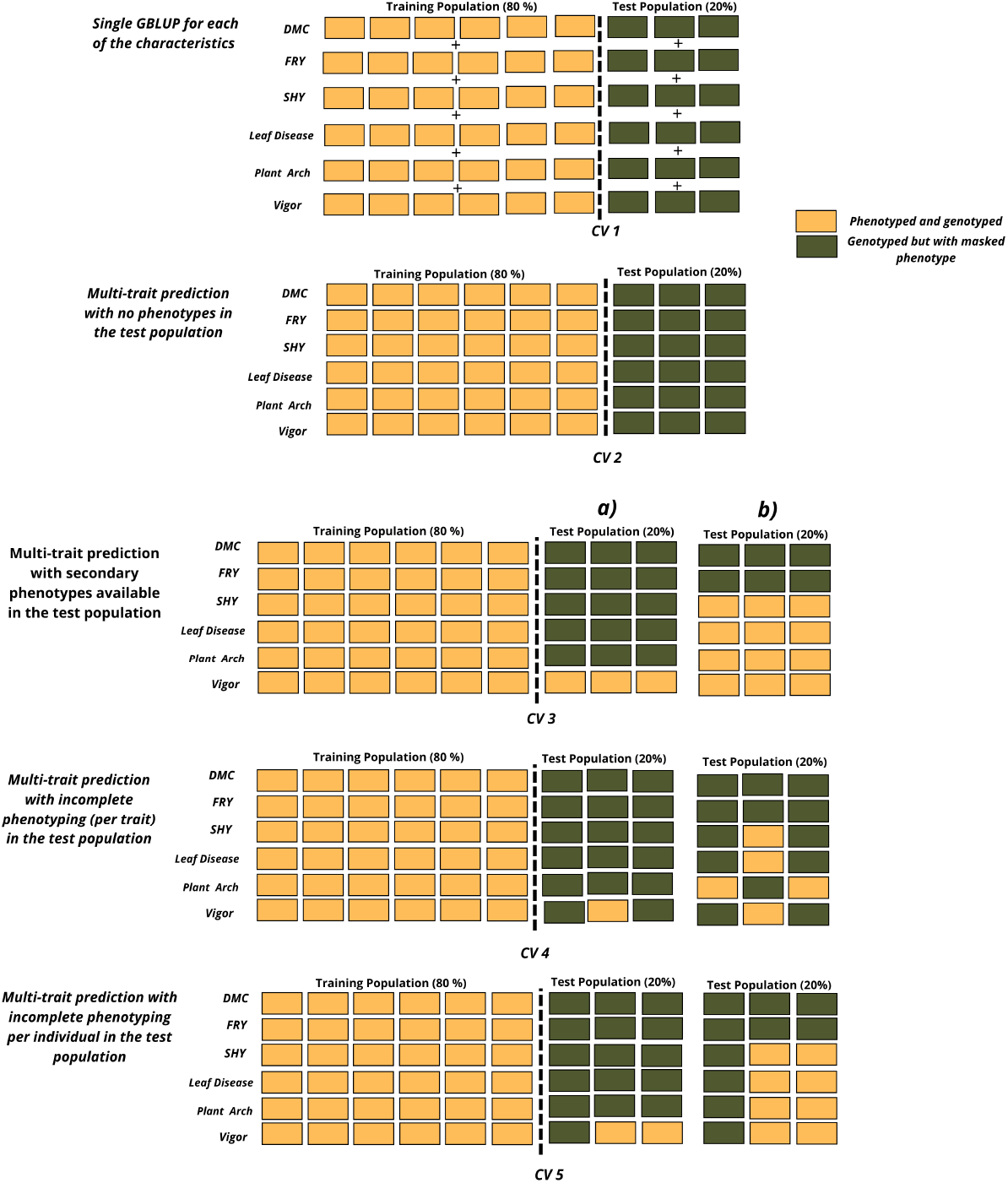
Representation of the five cross-validation (*CV*) scenarios used in univariate and multivariate genomic prediction models. *CV* 1 represents the single-trait model, which predicts each trait independently, disregarding correlations among traits. *CV* 2 – *CV* 5 correspond to multi-trait models, differing in the extent of phenotypic information available in the testing population. In *CV* 2, all six traits are predicted simultaneously using genotypic information only, without auxiliary phenotypes in the test set. In *CV* 3, secondary (auxiliary) traits are incorporated to jointly predict primary traits such as dry matter content and fresh root yield, leveraging inter-trait correlations to improve prediction accuracy. *CV* 4 and *CV* 5 follow the same multi-trait framework as *CV* 3 but simulate varying degrees of missing phenotypic information for the auxiliary traits: in *CV* 4, phenotypes are missing for specific traits within individuals, while in *CV* 5, they are missing for specific individuals within the population. As illustrative examples, panel (a) represents a model using a single auxiliary trait (Vigor), whereas panel (b) shows a model combining multiple auxiliary traits. *Abbreviations*: DMC – dry matter content; FRY – fresh root yield; SHY – shoot yield; Pl.Arch – plant architecture; Vigor – plant vigor at 1.5 months; Le.Dis – leaf disease severity.

By applying this CV across different trait combinations and data-missing scenarios, we quantified predictive ability under both ST and MT models. We emphasize that these CV schemes are designed to mimic breeder decisions: predicting performance of new, unphenotyped clones based solely on genotype and possibly auxiliary trait information. We also consider CV variations that simulate missingness by trait or clone to reflect sparse or partial phenotyping in real breeding programs.

CV 1 – Single-trait prediction (ST-GBLUP baseline): Each phenotypic trait was modeled independently, without considering correlations among traits. This baseline provides a reference for evaluating gains from multi-trait modeling. It represents typical genomic selection applications when inter-trait relationships are unknown, weak, or irrelevant to selection goals.

CV 2 – Multi-trait prediction with no phenotypes in the test population: The training population contains complete phenotypic information for both primary and secondary traits, while the test population consists solely of genotyped clones without phenotypic data. This setup mimics breeding situations in which newly genotyped clones have yet to be phenotyped, often due to time, cost, or environmental constraints. Evaluating this scenario shows how well genomic information alone predicts trait performance when no phenotypic data are available for the testing set.

CV 3 – Multi-trait prediction with secondary phenotypes available in the test population: Both training and test populations are genotyped. The training set has full phenotypic data, whereas the test set has phenotypic records only for secondary (easily measurable) traits. This scenario leverages known genetic correlations among traits to improve predictions for key primary traits, such as fresh root yield and dry matter content, which are typically labor-Intensive and costly to measure. In breeding practice, this approach allows programs to strategically phenotype only a subset of traits, thereby reducing costs while maintaining predictive power.

CV 4 – Multi-trait prediction with incomplete phenotyping per trait in the test population: Here, all test population clones are genotyped, but phenotypic data for secondary traits are partially missing across traits; some traits are measured on certain clones but missing on others. This design evaluates the model’s ability to borrow information across correlated traits to impute missing phenotypes, providing insights into how multi-trait models can mitigate information loss under partial phenotyping, a common limitation in large-scale field trials.

CV 5 – Multi-trait prediction with incomplete phenotyping per individual in the test population: In this configuration, only a subset of individuals in the test population is phenotyped for secondary traits, whereas the training set remains fully phenotyped. This setup simulates realistic resource constraints in cassava breeding pipelines, where only a portion of the population is phenotyped due to time, cost, or space constraints. The goal is to assess how predictive accuracy degrades or remains stable when secondary trait information is restricted to fewer individuals.

For CV 4 and CV 5, additional analyses were conducted by simulating 25%, 50%, and 75% missingness in the phenotypic data within both the training and testing subsets. Specify that these CVs were analyzed separately to clarify their distinction. These simulations enable the estimation of the minimum phenotyping intensity required to maintain satisfactory prediction accuracy.

**Table 1.** Combinations of primary and secondary traits across prediction scenarios (*CV* 3 to *CV* 5). In 13 of the 15 trait combinations, fresh root yield (FRY) and dry matter content (DMC) were jointly predicted, with an additional secondary trait selected based on its phenotypic Pearson correlation with both target traits. In the last two combinations, either FRY or DMC was individually defined as the primary trait, while the remaining traits were included as secondary traits in the multi-trait prediction model.

| Primary trait | Secondary trait |
| --- | --- |
| DMC+FRY+SHY+Pl.Arc+Leaf Disease | Vigor |
| DMC+FRY+Pl.Arc+Leaf Disease+Vigor | SHY |
| DMC+FRY+SHY+Leaf Disease+Vigor | Pl.Arc |
| DMC+FRY+SHY+Pl.Arc+Vigor | Leaf Disease |
| DMC+FRY+Pl.Arc+Leaf Disease | Vigor+SHY |
| DMC+FRY+SHY+Leaf Disease | Vigor+Pl.Arc |
| DMC+FRY+SHY+Pl.Arc | Vigor+Leaf Disease |
| DMC+FRY+Leaf Disease+Vigor | SHY+Pl.Arc |
| DMC+FRY+Pl.Arc+Vigor | SHY+Leaf Disease |
| DMC+FRY+SHY+Vigor | Leaf Disease+Pl.Arc |
| DMC+FRY+Vigor | SHY+Leaf Disease+Pl.Arc |
| DMC+FRY+SHY | Pl.Arc+Leaf Disease+Vigor |
| DMC+ FRY | SHY+Leaf Disease+Pl.Arc+Vigor |
| DMC | FRY+SHY+Leaf Disease+Pl.Arc+Vigor |
| FRY | DMC+SHY+Leaf Disease+Pl.Arc+Vigor |

### Evaluation metrics, coincidence analysis, and genetic gain

To assess the predictive performance and practical relevance of the genomic prediction models, we employed complementary evaluation metrics focusing on accuracy, precision, model consistency, and expected response to selection.

Model performance was primarily quantified through predictive ability (PA), defined as the Pearson correlation coefficient between the predicted GEBVs and the observed phenotypic BLUEs in the validation set. This metric quantifies the agreement between genomic predictions and adjusted phenotypic observations (Wolfe, Del Carpio, et al., 2017; Crossa et al., 2017).

In addition, the Root Mean Square Error (RMSE) was calculated to quantify the deviation between observed *y*_*i*_ and predicted *ŷ*_*i*_ values, providing a complementary measure of model precision :

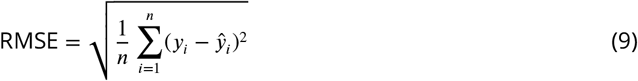

where *n* represents the number of observations, *y*_*i*_ is the observed value, and *ŷ*_*i*_ is the predicted value. Lower RMSE values indicate greater prediction accuracy and higher model stability across validation replicates.

To facilitate comparison among the different cross-validation scenarios, the relative change in PA and RMSE values obtained for each multivariate configuration (CV2–CV5) was expressed as the percentage improvement over the baseline single-trait model (CV1), using the following formula:

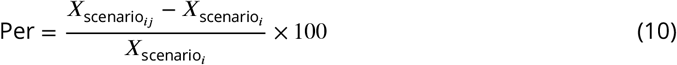

where *P er* represent the percentage variation for a given metric, 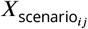 is the value of PA or RMSE obtained for the *j*_*th*_ combination of traits in the *i*_*th*_ CV scenario, and 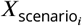 is the corresponding value for the univariate model (CV1).

The consistency of clone ranking across models was assessed using Cohen’s Kappa coefficient (*k*) (Cohen, 1960), which measures agreement beyond random expectation between the sets of top-ranked clones selected by the different genomic prediction models. For this analysis, the top-performing clones (10% selection intensity) based on GEBVs from the ST-GBLUP model (CV1) were compared with those identified by MT-GBLUP under the multi-trait scenarios (CV2–CV5). High *k* values indicate strong concordance in clone selection, suggesting that multi-trait models not only improve predictive accuracy but also preserve selection consistency relative to traditional univariate models.

To quantify the potential impact of improved predictive models on breeding progress, the response to selection (*RS*) was computed as:

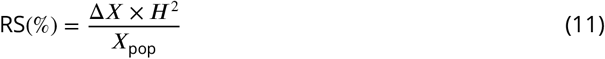

where Δ*X* is the selection differential (the difference between the mean of selected individuals and the overall population mean), *H*^2^ is the broad-sense heritability of the trait, and *X*_*pop*_ represents the population mean.

The expected genetic gain per cycle Δ*G* was also estimated according to Hayes et al. (2013):

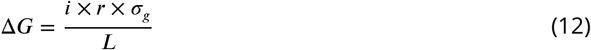

where *i* is the selection intensity (set to 10%), 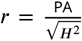 is the model accuracy, computed as is the standard deviation of the genotypic variance, and *L* represents the generation interval, assumed to be six years for cassava (two years for crossing and four years for clonal evaluation).

## Results

### Heritability estimates and trait correlations

The estimates of broad-sense heritability (*H*^2^) for the six evaluated traits ranged from low to moderate (Table 2). Dry matter content and fresh root yield exhibited moderate heritabilities (0.591 and 0.479, respectively), whereas leaf disease severity and plant vigor showed much lower *H*^2^ (0.108 and 0.202, respectively). The remaining traits, plant architecture and shoot yield, displayed intermediate heritabilities (0.308 and 0.361, respectively). When examining narrow-sense genomic heritability (*h*^2^), we observed slightly different orderings. Plant architecture, plant vigor, and leaf disease severity had a higher genomic *h*^2^, suggesting that additive genetic variance captured via SNPs is relatively higher for architecture than non-additive or residual components. The *h*^2^ estimates for dry matter content (0.498), fresh root yield (0.412), and shoot yield (0.341) were very similar to their corresponding *H*^2^ (Table 2).

**Table 2.** Broad sense heritability (*H*^2^) and Narrow-Sense genomic heritability (*h*^2^) for 6 traits evaluated along 1.078 clones in 8 environments of the brazilian cassava germplasm of EMBRAPA Cassava and Fruits, Cruz das Almas-BA,Brazil.

| Trait | $H^2$ | $h^2$ |
| --- | --- | --- |
| DMC | 0.591 | 0.498 |
| FRY | 0.479 | 0.412 |
| SHY | 0.361 | 0.341 |
| Pl.Arc | 0.308 | 0.462 |
| Leaf Disease | 0.108 | 0.197 |
| Vigor | 0.202 | 0.326 |

The phenotypic correlations (*r*_*p*_) among traits were generally low to moderate in magnitude, but mostly statistically significant (Figure 2). Traits that showed stronger phenotypic correlation often also displayed stronger genetic correlations (*r*_*g*_). For example, shoot yield and plant vigor exhibited *r*_*p*_ of 0.48, and their genetic correlation (0.64) was even higher, highlighting their potential utility as auxiliary traits in multivariate prediction. In contrast, leaf disease severity and plant architecture tended to show weak *r*_*p*_ and *r*_*g*_ with the other traits. For plant architecture, weak but significant positive *r*_*p*_ with fresh root yield (0.15) and shoot yield (0.12) were observed, whereas correlations with dry matter content (−0.018) and leaf disease severity (0.007) were non-significant. For leaf disease severity, only the correlation with dry matter content was positive and significant (0.06), though its magnitude was small; its *r*_*g*_ with dry matter content exceeded the *r*_*p*_, suggesting some underlying genetic association masked by environmental noise.

**Figure 2.**
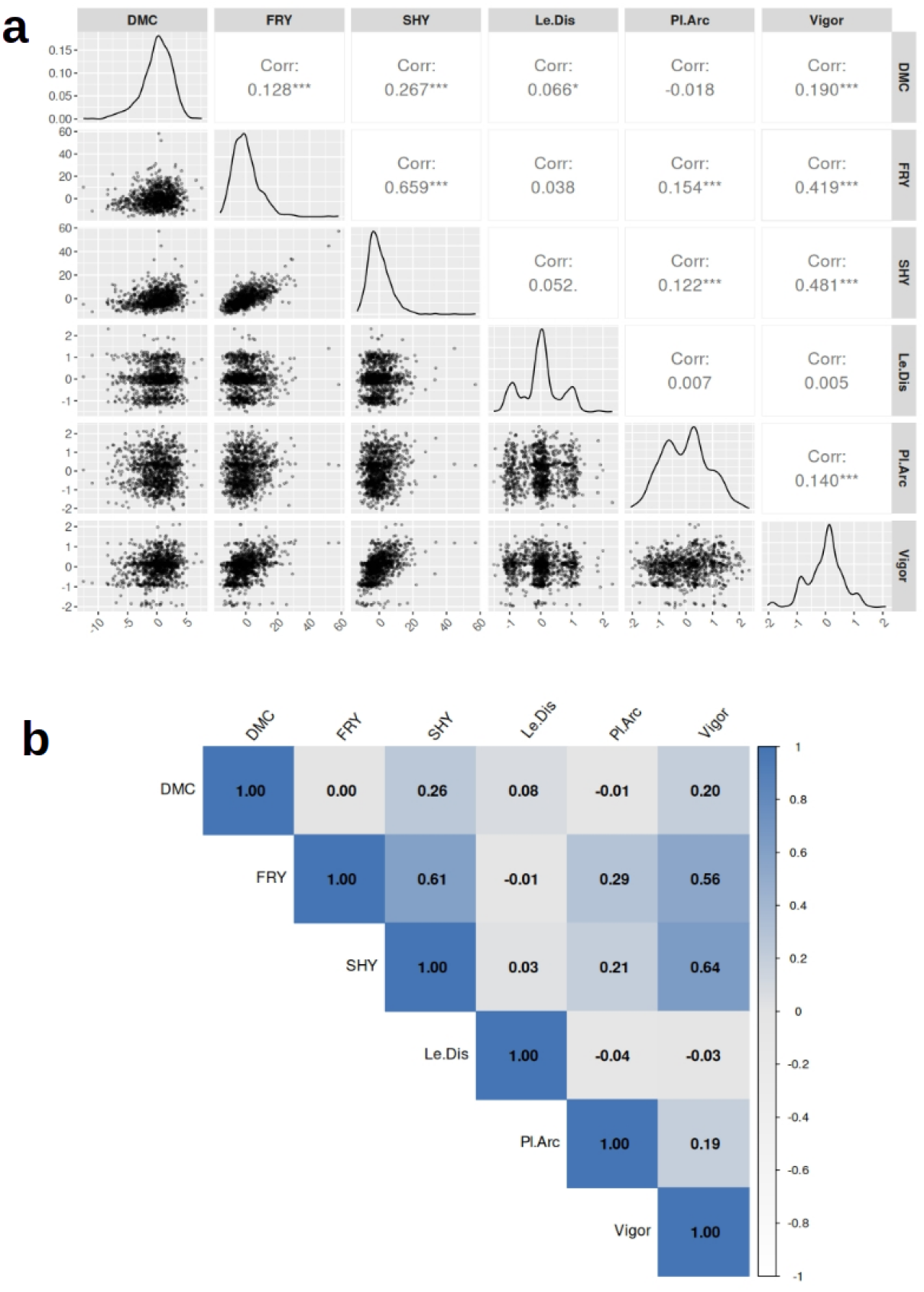
Phenotypic (*r*_*p*_) and genetic correlations (*r*_*g*_) among cassava traits. (a) Pairwise scatterplots, trait distributions, and Pearson phenotypic correlations among the BLUEs derived from a multi-environment trial analysis of 1,078 cassava clones. Diagonal panels show trait density distributions, lower panels display pairwise scatterplots, and upper panels present Pearson correlation coefficients with significance levels (*\* p* < 0.05, *\*\* p* < 0.01, *\*\*\* p* < 0.001). (b) Heatmap of genetic correlations among the evaluated traits, with color intensity representing the magnitude and direction of the correlations.Darker blue tones denote stronger positive correlations, darker white tones indicate stronger negative correlations, and intermediate gray shades represent moderate relationships.*Abbreviations*: DMC – dry matter content; SHY – shoot yield; Pl.Arc – plant architecture; FRY – fresh root yield; Vigor – plant vigor at 1.5 months; Le.Dis – leaf disease severity.

For dry matter content, *r*_*p*_ with other traits was generally low, with values of 0.26 with shoot yield, 0.19 with plant vigor, and 0.12 with fresh root yield, indicating that this trait is largely independent of yield-related phenotypes. In contrast, fresh root yield exhibited a broader and stronger correlation network, showing strong associations with shoot yield (*r*_*p*_ = 0.65) and plant vigor (*r*_*p*_ = 0.41), while correlations with the remaining traits were comparatively weak.

### Predictive performance between univariate (CV1) and multivariate (CV2) models

In the CV1 scenario, which considered the univariate prediction of each trait independently, predictive ability varied across traits, with the highest accuracy observed for dry matter content (0.50) and the lowest for leaf disease severity (0.13). For the remaining traits, fresh root yield, shoot yield, plant architecture, and vigor, predictive abilities were 0.45, 0.43, 0.46, and 0.42, respectively (Figures 3 and 6).

**Figure 3.**
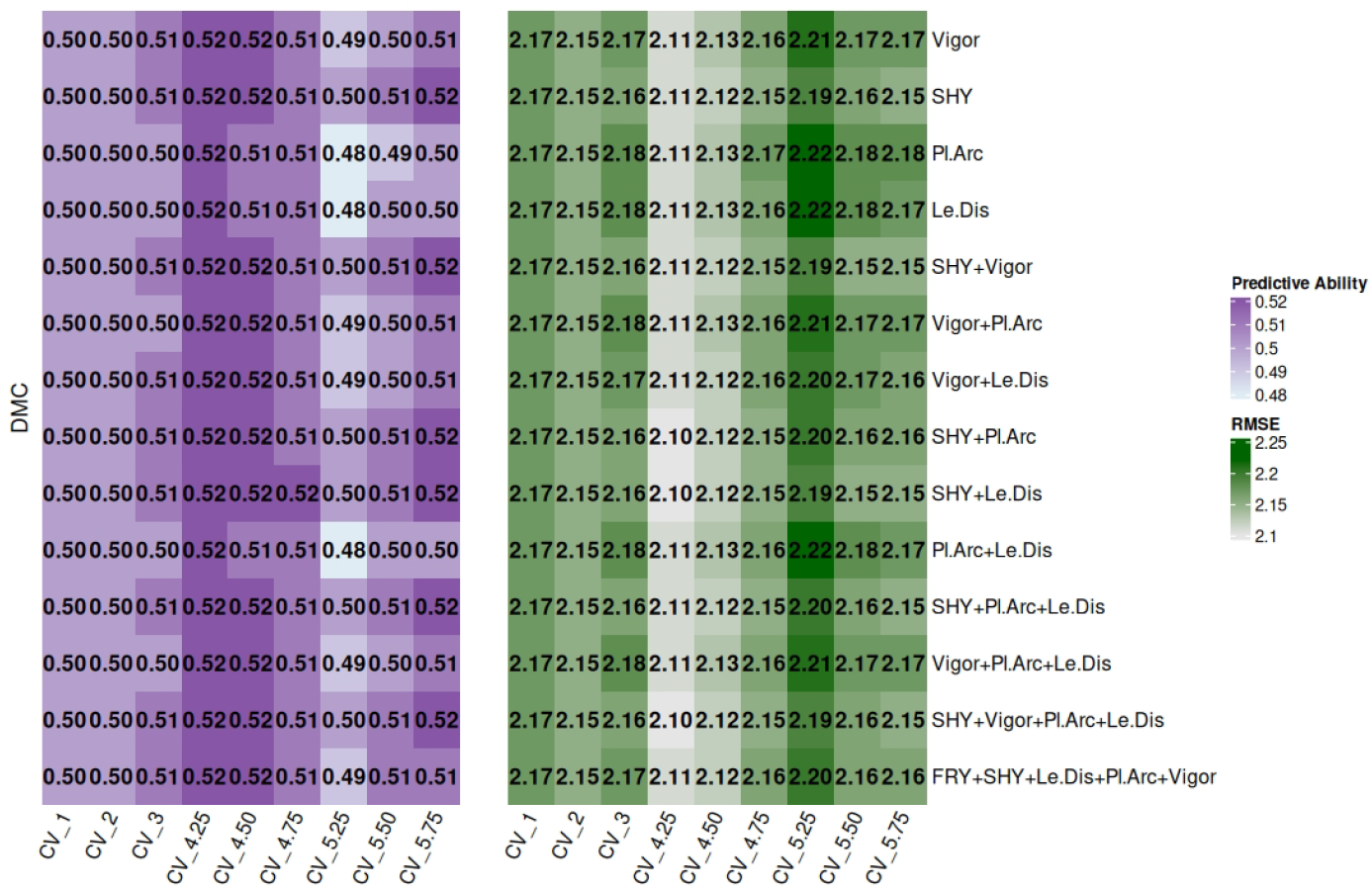
Comparison of predictive ability and the RMSE for dry matter content (DMC) prediction across different cross-validation scenarios: univariate model (CV1), multivariate prediction including all six traits (CV2), and multivariate models with incomplete phenotyping by trait (CV4) or by clone (CV5). For CV4 and CV5, progressive levels of phenotypic data absence (25%, 50%, and 75%) were simulated to assess model robustness under increasing data sparsity. *Abbreviations*: SHY – shoot yield; Pl.Arc – plant architecture; FRY – fresh root yield; Vigor – plant vigor at 1.5 months; Le.Dis – leaf disease severity.

In the CV2 scenario, where all six traits were jointly predicted using a fully multivariate model, the average predictive performance remained highly comparable to that of the univariate model. The corresponding accuracies were 0.50 (dry matter content), 0.46 (fresh root yield), 0.43 (shoot yield), 0.13 (leaf disease severity), 0.46 (plant architecture), and 0.42 (plant vigor). Similarly, changes in RMSE were modest, with fluctuations below ±2% relative to the univariate model. Thus, the univariate and fully multivariate approaches displayed nearly equivalent predictive performance across all evaluated traits.

### Predictive ability with highly correlated auxiliary traits (CV3)

In CV3, we evaluated the impact of incorporating secondary (auxiliary) traits into the joint prediction of dry matter content from the primary traits and fresh root yield, as well as their isolated predictions. Compared to CV2, which showed no substantial gains over CV1 (increases < 2%), CV3 revealed better improvements in both predictive ability and RMSE when correlated auxiliary traits were included. Traits exhibiting positive phenotypic correlations with dry matter content and fresh root yield, when used individually or in combination, produced performance gains, confirming the potential of trait-assisted multivariate prediction for complex agronomic traits. For dry matter content, predictive ability increased modestly, remaining similar to CV1 in some cases but improving by up to +2% in others. In contrast, fresh root yield showed high gains, with increases ranging from +2.22% to +44.44%. Correspondingly, reductions in RMSE were minor for dry matter content (up to −0.46%) but pronounced for fresh root yield (from −0.69% to −13.61%).

Among individual secondary traits, shoot yield emerged as the most informative, yielding the largest improvements in joint prediction: +2% for dry matter content (accuracy = 0.51) and +42.22% for fresh root yield (accuracy = 0.64). For two-trait combinations, shoot yield + plant vigor, shoot yield + plant architecture, and shoot yield + leaf disease severity stood out, with the shoot yield + leaf disease severity pair providing the best balance of gains: increasing predictive ability by +2% for dry matter content (0.51) and +44.44% for fresh root yield (0.65) while reducing RMSE by −0.46% for dry matter content (2.16) and −13.06% for fresh root yield (6.26) (Figures 3 and 4).

**Figure 4.**
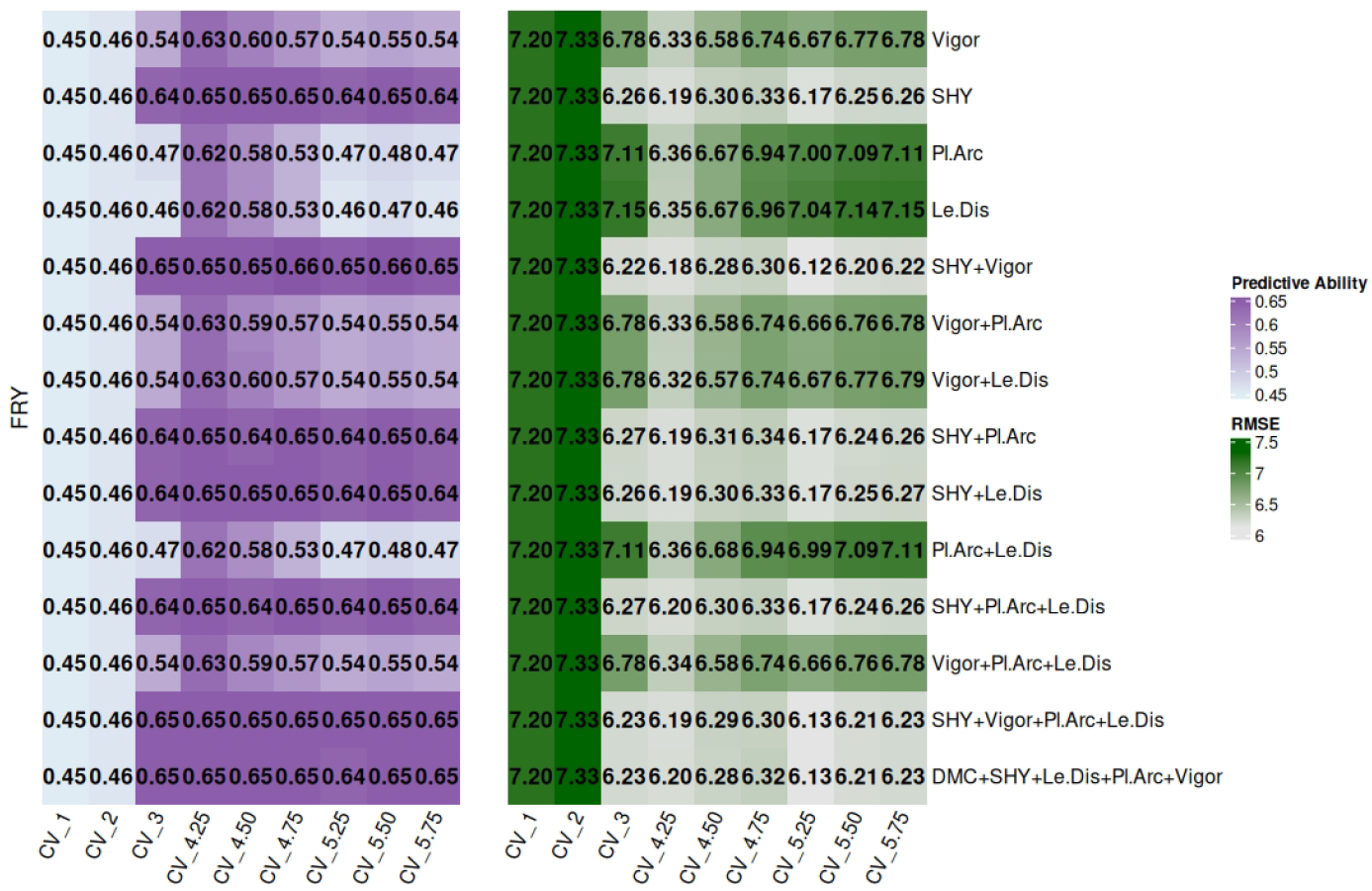
Comparison of predictive ability and the RMSE for fresh root yield (FRY) prediction across different cross-validation scenarios: univariate model (CV1), multivariate prediction including all six traits (CV2), and multivariate models with incomplete phenotyping by trait (CV4) or by clone (CV5). For CV4 and CV5, progressive levels of phenotypic data absence (25%, 50%, and 75%) were simulated to evaluate model robustness under increasing data sparsity. *Abbreviations*: DMC – dry matter content; SHY – shoot yield; Pl.Arc – plant architecture; Vigor – plant vigor at 1.5 months; Le.Dis – leaf disease severity.

When three auxiliary traits were combined, shoot yield + plant architecture + leaf disease severity yielded the greatest benefits, achieving +2% and +44.44% gains for dry matter content and fresh root yield, respectively, accompanied by RMSE reductions of −0.46% (2.16) and −13.47% (6.23). The inclusion of four traits produced similarly consistent improvements (dry matter content: +2%, fresh root yield: +44.44%).

Finally, when all secondary traits were included (for dry matter content: fresh root yield + shoot yield + leaf disease severity + plant architecture + plant vigor; and for fresh root yield: dry matter content + shoot yield + leaf disease severity + plant architecture + plant vigor), effects differed between the two focal traits. For dry matter content, predictive ability improved modestly by 2%, with no meaningful reduction in RMSE. Conversely, fresh root yield benefited substantially, exhibiting a 44.44% increase in predictive ability (0.65) and a 13.47% reduction in RMSE (6.23) (Figures 5 and 6) When a trait was used as an auxiliary for predicting fresh root yield and dry matter content, all other evaluated traits were simultaneously included in the prediction process. For example, when plant vigor was used as an auxiliary trait, the prediction included dry matter content, fresh root yield, shoot yield, plant architecture, and leaf disease severity. Importantly, the combinations that maximized predictive ability and minimized RMSE for fresh root yield and dry matter content did not always coincide with the optimal combinations for the other traits.

**Figure 5.**
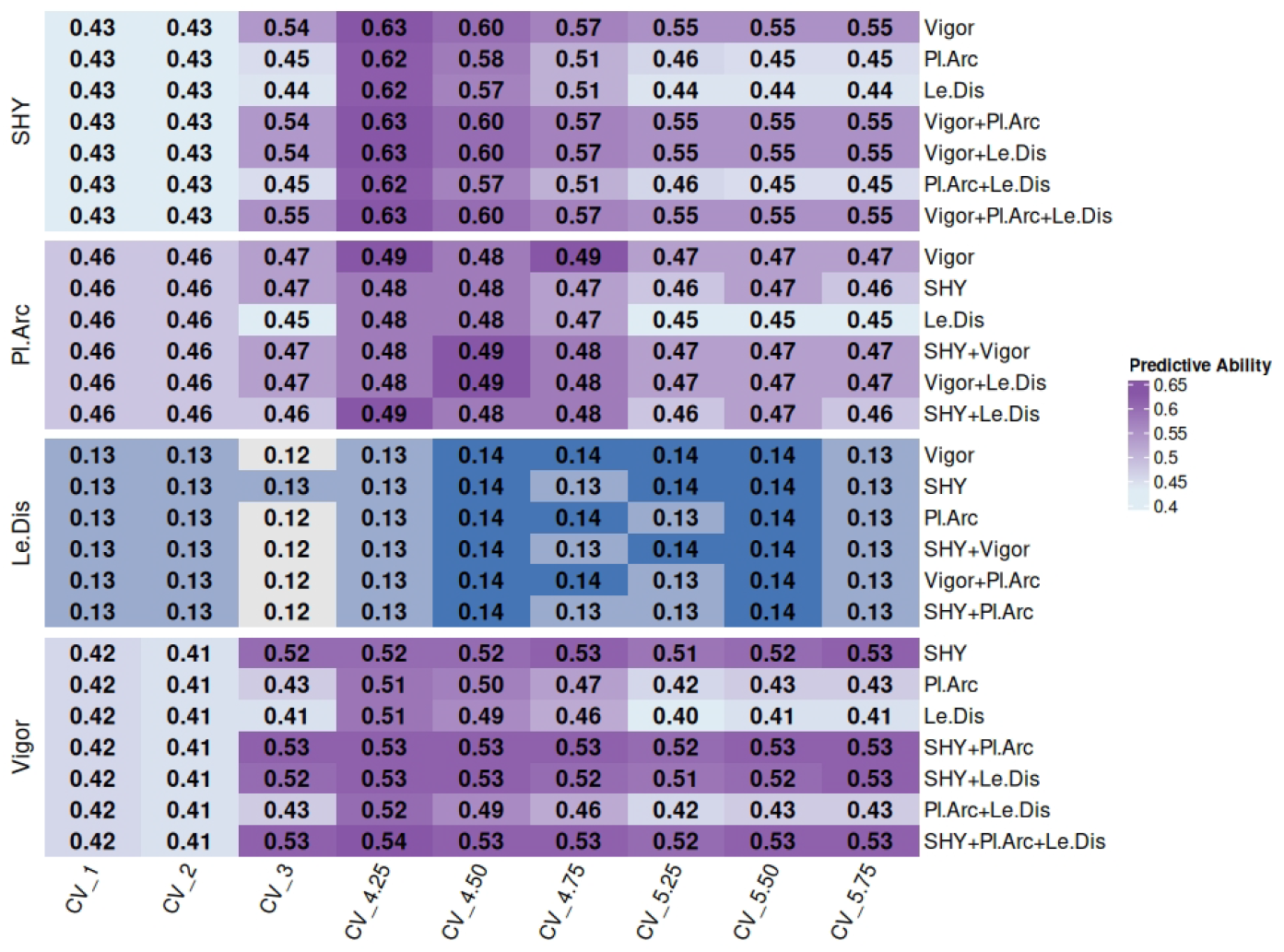
Predictive ability for shoot yield (SHY), leaf disease severity (Le.Dis), plant architecture (Pl.Arch), and plant vigor (Vigor) across different cross-validation scenarios: univariate model (CV1), multivariate joint prediction of six traits (CV2), and multivariate scenarios with incomplete phenotyping per trait (CV4) or per clone (CV5). For CVs 4 and 5, simulations illustrate progressively increasing levels of complexity due to 25%, 50%, and 75% missing phenotypic data. DMC – dry matter content; FRY – fresh root yield; Le.Dis – leaf disease severity.

**Figure 6.**
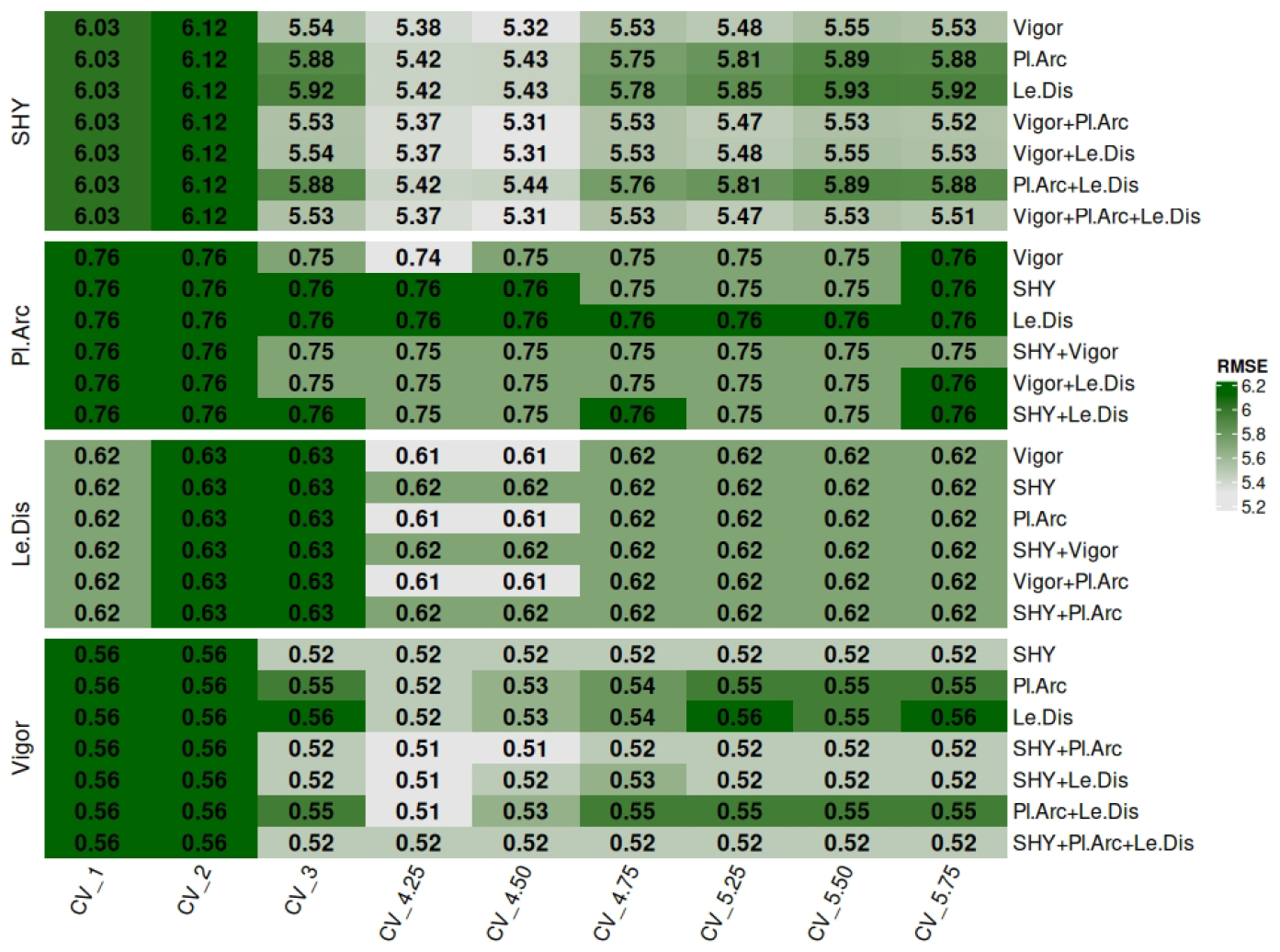
RMSE for shoot yield (SHY), leaf disease severity (Le.Dis), plant architecture (Pl.Arch), and plant vigor (Vigor) across different cross-validation scenarios: univariate model (CV1), multivariate joint prediction of six traits (CV2), and multivariate scenarios with incomplete phenotyping per trait (CV4) or per clone (CV5). For CVs 4 and 5, simulations depict progressively increasing complexity associated with 25%, 50%, and 75% missing phenotypic data. DMC – dry matter content; FRY – fresh root yield; Le.Dis – leaf disease severity.

Shoot yield predictions benefited from the inclusion of plant vigor, either individually or in combination with plant architecture, leaf disease severity, or all three traits simultaneously. The greatest improvement was observed with the combination of plant vigor + plant architecture + leaf disease severity, resulting in a 27.91% increase in predictive ability (0.55) and an 8.29% reduction in RMSE (5.53). For leaf disease severity, the most informative auxiliary trait was shoot yield. However, predictive accuracy remained at the same level as the univariate model (0.13), and RMSE increased slightly by 1.61%, regardless of the combination tested (Figures 5 and 6).

Regarding plant architecture, the best results were obtained using plant vigor, either alone or combined with leaf disease severity or shoot yield. All these combinations led to a 2.17% increase in predictive ability (0.47) and a 1.32% reduction in RMSE (0.75). For plant vigor, the largest gains were observed when combined with shoot yield, either alone or together with plant architecture and/or leaf disease severity. Among these combinations, shoot yield + plant architecture and shoot yield + plant architecture + leaf disease severity were the most effective, producing a 26.19% increase in predictive ability (0.53) and a 7.14% reduction in RMSE (0.52) (Figures 5 and 6).

### Predictive ability under balanced and unbalanced phenotypic data scenarios

Cross-validation scenarios 4 and 5 (CV4 and CV5) were designed to evaluate model performance under different patterns and degrees of missing phenotypic data, representing practical constraints in breeding programs. CV4 simulated unbalanced data with missing values distributed by trait, whereas CV5 simulated balanced missing data with missing values distributed by individual clones. Each scenario was further stratified to include 25%, 50%, and 75% missing data, allowing a systematic assessment of how varying data availability affects predictive ability and RMSE. The analysis also evaluated the stabilizing role of secondary traits in multivariate prediction models.

For dry matter content and fresh root yield, the largest gains relative to the univariate model (CV1) and the multivariate CV2 were consistently achieved when shoot yield was used as a secondary trait, either alone or in combination with plant vigor, plant architecture, and leaf disease severity. Combinations of two traits (shoot yield + plant vigor, shoot yield + plant architecture, shoot yield + leaf disease severity), three traits (shoot yield + plant architecture + leaf disease severity), or four traits (shoot yield + plant architecture + leaf disease severity + plant vigor) provided the most stable improvements. These combinations, previously highlighted in CV3, confirmed their robustness as auxiliary traits in maintaining high PA and low RMSE across missing-data scenarios.

Compared with the CV3 scenario, both CV4 (missing data by trait) and CV5 (missing data by clone) enabled robust simultaneous prediction of fresh root yield and dry matter content, as assessed by predictive ability and RMSE. For dry matter content, gains were modest but consistent relative to CV1–CV3, with stability maintained when previously validated favorable combinations of secondary traits were applied. This stability persisted even under simulated missing-data conditions (25%, 50%, and 75%). Predictive ability for dry matter content remained stable across CV4 and CV5, with only minor variations among missing data proportions. Specifically, CV4 produced gains of 2–4% relative to univariate (CV1) and other multivariate models, with the largest improvement (4%, predictive ability 0.52) observed at 25% missing data. In CV5, slight reductions were noted in some combinations (up to −4%), but most scenarios demonstrated gains of 2–4% or performance comparable to CV1 (Figure 3).

For fresh root yield, predictive performance was more sensitive to the proportion of missing data. The most favorable trait combinations consistently produced gains surpassing those of univariate and CV2–CV3 models. Even less optimal combinations maintained predictive ability improvements with only minor RMSE changes, while the best combinations sustained substantial and consistent gains under high missing data levels (50% and 75%) in both CV4 and CV5 (Figure4). Overall, the highest predictive performance for fresh root yield and dry matter content was observed in CV4 with 25% missing data, with predictive ability ranging from 0.62 to 0.65, which decreased slightly at 50% and 75% missingness. In CV5, predictions were more stable, with the most favorable outcomes at 50% missing data and minimal fluctuations across different missing data levels, maintaining predictive ability values between 0.46 and 0.65 (Figures 3 and 4)

RMSE patterns differed between fresh root yield and dry matter content. For dry matter content, increasing missing data in CV4 led to small increases in RMSE, i.e., 2.10–2.11 at 25%, 2.12–2.13 at 50%, and 2.15–2.17 at 75% missing data. In CV5, the trend was reversed, with RMSE remaining close to CV1 values and showing minor fluctuations: lowest at 75% missing (2.15–2.16), slightly higher at 50% (2.16–2.18), and largest at 25% (2.18–2.22), representing a modest RMSE increase compared to other scenarios (Figures 3 and 4). For fresh root yield, both CV4 and CV5 reduced RMSE relative to CV1–CV3. In CV4, the greatest reductions occurred at 25% missing data (6.18–6.36), followed by 50% (6.30–6.67) and 75% (6.30–6.96). CV5 exhibited a similar trend, with RMSE values of 6.30–6.67 at 25%, 6.20–7.14 at 50%, and 6.22–7.11 at 75% missing data.

For the remaining traits, the combinations previously identified as favorable in CV3 continued to yield consistent gains in predictive ability and reductions in RMSE compared with the CV1–CV3 scenarios. For shoot yield, combinations involving plant vigor, plant architecture, and leaf disease severity consistently enhanced predictive ability and reduced RMSE, particularly under CV4 with 25% missing data. In CV5, these gains remained stable across varying levels of missing data, indicating robustness of the predictive model (Figures 5 and 6)

For leaf disease severity and plant architecture, the most favorable auxiliary combinations included either shoot yield or plant vigor alongside other traits, with the shoot yield + plant vigor combination emerging as particularly advantageous for both traits. Under CV4, the highest predictive ability for leaf disease severity was achieved with 50% missing data, while at the extremes (25% and 75%), predictive ability remained comparable to that of CV1. RMSE reductions were modest, with values generally maintaining CV1 levels. In CV5, predictive ability and RMSE were largely unaffected by the proportion of missing data, remaining stable across all secondary trait combinations.

For plant architecture, combinations involving plant vigor or shoot yield + leaf disease severity produced the largest increases in predictive ability and corresponding reductions in RMSE under CV4. In CV5, gains were more modest but remained stable across missing-data proportions, showing minimal sensitivity to data imbalance.

For plant vigor, the combination of shoot yield and plant architecture proved most favorable, delivering consistent gains in predictive ability in both CV4 and CV5, with little variation across missing-data levels. RMSE reductions were more pronounced in CV4, decreasing progressively as the proportion of missing data increased. In C increasedV5, RMSE remained stable, comparable to CV1 values, indicating robust model performance under variable data completeness scenarios (Figures 5 and 6).

### Multi-trait genomic prediction enhances selection response

We further evaluated selection response and estimated genetic gains across CV scenarios, focusing on auxiliary-trait combinations that maximized predictive ability while minimizing RMSE for fresh root yield and dry matter content. In CV2, the joint multivariate model across all six traits outperformed the univariate model (CV1) for most traits. Exceptions were dry matter content (4.30 vs. 4.20 in CV1), fresh root yield (27.47 vs. 27.22 in CV1), and plant architecture (10.17 vs. 9.60 in CV1). Across CV3–CV5, gains and modest reductions in selection response occurred depending on the trait and auxiliary-trait combination. For dry matter content, gains ranged from 0.51% (plant vigor + plant architecture + leaf disease severity) to 1.48% (leaf disease severity alone), mainly in CV3 and CV5, where improvements spanned 0.51–1.83% and were associated with leaf disease severity. In CV4, minor reductions ranged from −1.19% (shoot yield + plant architecture with 50% missing data) to −4.19% (plant vigor with 75% missing data).

For fresh root yield, gains ranged from 0.20% (plant architecture + leaf disease severity, CV4, 50% missing data) to 2.67% (plant vigor + plant architecture + leaf disease severity, CV4, 25% missing data), whereas modest decreases occurred in CV3 and CV5, ranging from −0.97% to −2.78%. Shoot yield consistently benefited from auxiliary trait inclusion. Gains varied from 0.17% (plant architecture + leaf disease severity, CV4, 50% missing data) to 3.60% (plant architecture, CV4, 75% missing data), with the largest increase observed in CV2 (5.56%, from 13.90 to 14.68).

For plant architecture, small reductions predominated, ranging from −0.40% (shoot yield + plant vigor + plant architecture + leaf disease severity, CV3) to −4.45% (leaf disease severity, CV4, 50% missing data). In contrast, plant vigor and leaf disease severity exhibited consistent gains across CV3–CV5. Plant vigor increased 6.97%–10.78%, and leaf disease severity increased 3.18%–9.72%, depending on the auxiliary trait combination and missing data scenario.

### Multi-trait genomic prediction enhances genetic gains

Analysis of genetic gains showed that the multi-trait genomic prediction model substantially improved performance, particularly when highly correlated traits were included, as observed in CV3–CV5. These gains were evident for individual traits and their combinations. Even with moderate correlations, such as plant vigor alone or combined with plant architecture and leaf disease severity for dry matter content, small but meaningful gains were detected. The stability of these responses across validation scenarios and levels of missing data indicates that secondary traits can amplify selection gains while maintaining predictive robustness under practical breeding conditions.

In CV2, compared to the univariate model, gains were modest for dry matter content, fresh root yield, leaf disease severity, and plant vigor, whereas shoot yield and plant architecture showed slight reductions of −0.86% and −2.16%, respectively. In contrast, the inclusion of highly correlated traits in CVs 3, 4, and 5 led to pronounced improvements for most traits. For dry matter content, increases were generally close to the univariate baseline or ranged from 2% to 4% across the CV3 and CV4–5 scenarios with 25–75% missing data. Fresh root yield exhibited larger gains, ranging from 2.22% to 44.44%, consistent across CV3 and CV4–5 despite varying proportions of missing data. Shoot yield gains ranged from 21% to 42%, particularly in CV4 with highly correlated trait combinations, while CV5 showed only slight declines. For leaf disease severity, CV3 resulted in reductions of approximately −7.96% in certain combinations, whereas others matched or exceeded univariate performance. In CVs 4 and 5, gains were observed with 25% missing data, but higher levels of missingness (50% and 75%) produced moderate reductions. Plant architecture gains were more modest (2.17%–6.52%) and were particularly pronounced in CV4 and CV5 with 25% missing data. Plant Vigor stood out with consistent gains ranging from 9.52% to 26.19%, achieving the highest performance in CV4 and CV5 while maintaining stability across all missing-data levels.

### Consistency of selection: Kappa index between single-trait and multi-trait models

To evaluate the reliability of MT predictions for selection, the Kappa Index was calculated relative to the univariate model (CV1) for all auxiliary-trait combinations and validation scenarios. Overall, concordance was highest when auxiliary traits effectively improved prediction of the target trait. In CV2, Kappa values ranged from intermediate (0.67) to high (> 0.80), indicating strong agreement between the top 10% of clones selected under MT versus univariate models. CV3 showed similarly high concordance, except for some combinations involving leaf disease severity and plant architecture, which slightly decreased (< 0.70), while the remaining combinations stayed between 0.70 and 0.80. In CV4 and CV5, concordance was predominantly high (> 0.80), independent of missing-data levels, although leaf disease severity and plant architecture occasionally showed slightly lower values (0.63–0.76).

## Discussion

### Revisiting the baseline: situating univariate and simple multivariate predictions in cassava genomics

Genomic selection has transformed cassava breeding by accelerating selection gains and reducing reliance on extensive phenotyping cycles, particularly within the Brazilian cassava breeding program. Historically, genomic selection efforts have primarily used univariate GBLUP models to predict key agronomic and industrial traits such as dry matter content and fresh root yield, which are crucial for processing and industrial applications. These models have successfully delivered moderate-to-high predictive abilities, supporting their continued role in cassava improvement programs targeting medium- and long-term selection gains (Torres et al., 2019; Andrade, Sousa, Oliveira, et al., 2019; Andrade, Sousa, Wolfe, et al., 2022; Costa et al., 2024).

In this study, our baseline univariate GBLUP (CV1) achieved consistent, biologically meaningful predictive performance across traits, with intermediate predictive performance and low RMSE. This outcome corroborates prior findings for Brazilian germplasm, where predictive ability values for dry matter content ranged between 0.51–0.59 (Torres et al., 2019) and for fresh root yield between 0.43–0.50 (Andrade, Sousa, Oliveira, et al., 2019; Andrade, Sousa, Wolfe, et al., 2022), whereas Costa et al. (2024) reported slightly lower estimates (0.30 for dry matter content and 0.41 for fresh root yield). Predictive performance for shoot yield, plant vigor, and plant architecture was also within literature-reported ranges, with shoot yield outperforming Costa et al. (2024) (0.31) and approximating Torres et al. (2019) (0.48–0.50). Plant vigor estimates aligned with Okeke et al. (2017) (0.16–0.42), while plant architecture, characterized by relatively stable phenotypic expression (Sampaio Filho et al., 2023; Dos Santos et al., 2023), also exhibited positive predictive outcomes. The lowest predictive ability values were obtained for leaf disease severity (0.13), consistent with the large variation reported in disease-prediction studies for cassava brown streak disease (0.05–0.41) (Ozimati et al., 2018; Kayondo et al., 2018) and cassava mosaic disease (−0.02–0.59) (Okeke et al., 2017; Wolfe, Del Carpio, et al., 2017). These low accuracies are biologically plausible given the complex, polygenic nature of disease resistance and the strong environmental modulation of expression.

Overall, the baseline results confirm that univariate GBLUP remains a stable and reliable framework for marginal predictions of individual traits in cassava. However, its conceptual limitation is its inability to exploit cross-trait genetic and phenotypic covariances, thereby constraining predictive accuracy for low-heritability traits and limiting the efficiency of selection response. To address this, multivariate GBLUP models, which leverage inter-trait correlations, have emerged as a promising extension of univariate frameworks (Calus and Veerkamp, 2011; Lado et al., 2018; Bhatta et al., 2020; Dhakal et al., 2024).

In our study, CV2 represented the first multivariate scenario, involving simultaneous prediction of six phenotypic traits in a newly genotyped but unphenotyped population. The performance of MT-GBLUP under all-traits configuration was broadly equivalent to that of univariate GBLUP in both predictive ability and RMSE, with similar findings in other crops where weak inter-trait correlations limit multivariate gains (Anilkumar et al., 2025; Fernandes et al., 2018). This pattern is consistent with the correlation structure observed in Figure 2, where most phenotypic and genetic correlations among traits were weak to moderate, and occasionally negative. An exception was shoot yield, which showed strong positive correlations with several traits, particularly fresh root yield, but only a weak association with leaf disease severity. Although this relatively limited genetic covariance structure could reduce the potential advantage of MT-GBLUP in this configuration (Jia and Jannink, 2012), the simultaneous prediction of all six traits still proved beneficial.

Despite these constraints, the baseline results underscore the importance of dry matter content and fresh root yield within cassava improvement programs. Dry matter content directly affects processing quality and economic value through its influence on product yield (flour, starch, and other derivatives), while fresh root yield serves as a direct estimate of field productivity (Joaqui Barandica et al., 2016). These traits, though central to industrial and commercial goals, are costly and time-consuming to phenotype, making them ideal targets for predictive genomics, especially through trait architectures that leverage auxiliary information from correlated, inexpensive traits.

### Leveraging correlated traits to enhance prediction accuracy and RMSE reduction

Targeted multivariate designs (CV3) were implemented to exploit strong inter-trait correlations, particularly between fresh root yield and auxiliary traits such as shoot yield, plant vigor, and leaf disease severity. Under this setup, MT-GBLUP delivered clear gains in predictive ability and reductions in RMSE for both dry matter content and fresh root yield, confirming the theoretical advantage of incorporating correlated secondary traits (Bhatta et al., 2020; Lado et al., 2018; Anilkumar et al., 2025; Dhakal et al., 2024; Fernandes et al., 2018). The trait shoot yield emerged as the most promising auxiliary trait for improving prediction of both fresh root yield and dry matter content, especially when combined with plant vigor and leaf disease severity. This result aligns with the prediction obtained by Jia and Jannink (2012) that increasing genetic correlations among traits boosts multivariate predictive accuracy, an effect most pronounced for traits with low heritability.

The success of shoot yield as an auxiliary trait may be related to the fact that shoot biomass and vigor represent aboveground indicators of photosynthetic capacity and source strength, both of which are closely linked to root bulking and assimilate partitioning. Therefore, shoot yield, being a relatively easy, fast, and early trait to phenotype, serves as a biologically and operationally efficient proxy for fresh root yield, enhancing the overall informational content of genomic predictions. Similar relationships between biomass and yield have been explored in cereals and forages, where early-stage correlated phenotypes have improved yield prediction through multivariate (MT) models (Arojju et al., 2020; Velazco et al., 2019; Fradgley et al., 2023).

Conversely, traits such as leaf disease severity and plant architecture, while quite inexpensive to measure, offered weaker or inconsistent improvements in prediction due to their low or negative correlations with the primary traits. This reinforces the necessity of strategic auxiliary-trait selection, prioritizing traits with demonstrably favorable genetic covariance structures, to avoid dilution of predictive signal and to maximize resource-use efficiency. The improved performance in CV3 relative to CV1 and CV2 thus demonstrates that when auxiliary traits are carefully selected based on correlation and biological relevance, MT-GBLUP effectively increases predictive reliability, particularly for complex, polygenic traits with moderate-to-low heritability such as fresh root yield.

### The role of missing data and sparse phenotyping designs

The CV4 and CV5 scenarios extended the analysis to breeding-relevant contexts characterized by incomplete phenotypic datasets, a common challenge in cassava improvement due to METs, logistical constraints, and resource limitations. These simulations tested the robustness of MT-GBLUP under varying proportions of missing data and different missingness structures (by trait in CV4, by clone in CV5). The results revealed that multivariate models maintained or even improved predictive ability relative to univariate baselines, despite increasing proportions of missing data, demonstrating their robustness and practical utility under real-world conditions.

In CV4, the best performance occurred when 25% of the phenotypic data were missing, with prediction accuracy decreasing gradually at higher levels of missingness (50–75%). Conversely, CV5 showed a counterintuitive pattern, where accuracy for dry matter content and fresh root yield peaked at 75% missing data. These outcomes indicate that partial but strategically distributed phenotyping, where auxiliary traits remain well represented across genotypes, can sustain or even enhance model performance. This finding aligns with previous reports from cereals and legumes demonstrating that sparse phenotyping designs, when optimized for trait covariance and genetic connectedness, maximize prediction accuracy while minimizing cost (Lado et al., 2018; Gaire et al., 2022; Lubanga et al., 2025).

In practice, this implies that cassava breeding programs can reduce the phenotyping burden without sacrificing predictive power by leveraging correlated traits and well-designed sparse testing schemes. This approach is particularly beneficial for labor-intensive traits such as dry matter content and fresh root yield, which require destructive sampling and extensive fieldwork. Moreover, as shown in Gaire et al. (2022), integrating optimization algorithms into phenotyping design can ensure that data collection maximizes information flow across genotypes and environments. Together, these findings position MT-GBLUP as a key operational tool for breeding programs facing budgetary and logistical limitations.

### Consistency and selection response under MT genomic prediction

One of the most striking results was the high concordance between the best-performing genotypes identified under univariate and multivariate models, with Kappa indices typically exceeding 0.80 across cross-validations, underscoring the stability of selection decisions. This robustness persisted even in the presence of missing phenotypic data, confirming the model’s practical resilience and its ability to preserve ranking accuracy under realistic breeding conditions.

The moderate reductions in concordance observed for leaf disease severity and plant architecture (Kappa = 0.63–0.76) can be attributed to their more complex genetic architecture and higher G×E interactions, which are known to introduce instability into selection responses (Costa et al., 2024; Andrade, Sousa, Oliveira, et al., 2019). Additionally, the high selection intensity (10%) likely accentuated small differences in predicted performance among top clones, as reported (Costa et al., 2024).

Beyond selection consistency, multivariate models outperformed univariate models in selection response and genetic gain per unit time, particularly for low-heritability or labor-intensive traits such as shoot yield, plant vigor, and leaf disease severity. These gains are quite similar to those observed in other clonally propagated crops, where MT-GBLUP has been shown to accelerate realized selection efficiency by exploiting correlated trait information (Ehoche et al., 2025; Matias et al., 2019; Bhatta et al., 2020). Importantly, these improvements persisted even when large proportions of phenotypic data were missing (CV4, CV5), reinforcing the feasibility of MT-GBLUP for continuous, cost-effective selection in cassava. Thus, these results demonstrate that MT-GBLUP not only maintains high selection stability but also enhances the rate of genetic gain per selection cycle, offering a path to shorten breeding cycles and achieve higher cumulative gains over time.

In addition to maintaining selection consistency, multivariate models outperformed univariate models in response to selection and genetic gain per unit of time, particularly for traits with low heritability or that are difficult and labor-intensive to measure, such as fresh root yield and dry matter content. In this study, these gains were primarily driven by a reduction in breeding cycle length and the ability to generate reliable predictions under sparse phenotyping scenarios, as demonstrated in CV4 and CV5. Similar advantages of MT-GBLUP have been reported in other clonally propagated crops, where multi-trait models improved selection efficiency by leveraging information from genetically correlated traits (Ehoche et al., 2025; Matias et al., 2019; Bhatta et al., 2020). Notably, these improvements were maintained even under high levels of missing phenotypic data (CV4 and CV5), reinforcing the potential of MT-GBLUP as a cost-effective strategy for continuous selection in cassava breeding. Overall, our results indicate that MT-GBLUP not only preserves selection stability but also enhances genetic gain per selection cycle, providing a practical approach to shorten breeding cycles and increase cumulative genetic gains over time.

### Suggestion for future model extensions

Recent developments in multi-trait machine learning (MT-ML) and deep learning offer promising avenues for modeling nonlinear relationships, epistasis, and complex genotype–environment interactions. Studies in maize, wheat, and rice have shown that multi-kernel ensembles and deep neural networks can outperform linear GBLUP when sufficient training data and trait correlation structures are available (Montesinos-López et al., 2018; Fradgley et al., 2023). However, such approaches also demand rigorous cross-validation, large and heterogeneous training datasets, and careful model regularization to avoid overfitting, requirements that remain challenging in cassava breeding contexts with limited phenotypic and environmental diversity.

Therefore, MT-ML approaches should be considered as complementary tools to traditional MT-GBLUP rather than replacements. When integrated judiciously, these methods may further enhance predictive resolution and accelerate selection gains for complex traits such as yield and disease resistance.

### Translational perspectives for integration into cassava breeding programs

Applying empirical findings from multi-trait genomic prediction to routine cassava breeding requires a pragmatic, evidence-based strategy, similar to approaches successfully implemented in other crops. Here, we integrate our results with transferable lessons from other crops and include steps for pilot testing, such as sparse phenotyping, high-throughput phenotyping (HTP) integration, and the progressive refinement of prediction models. Initial implementation should focus on targeted pilot studies, combined with economic validation, to compare MT and ST workflows in terms of candidate-ranking agreement (Kappa), predictive ability, and short-term genetic gains. Routine estimation of genetic correlations among traits should also be prioritized to identify auxiliary traits that are inexpensive to measure yet highly informative for prediction, as observed for shoot yield and plant vigor in our dataset. In addition, pilot studies should incorporate economic assessments, including phenotyping cost per plot, labor and time requirements, and projected annual genetic gain, since predefined cost-benefit criteria have been critical for the successful adoption of genomic prediction strategies in wheat and maize breeding programs (Lado et al., 2018; Matias et al., 2019).

HTP can supply the inexpensive, repeatable auxiliary signals MT models need, but only when those proxies show heritability and genetic correlation with focal traits. (Carvalho et al., 2022) image-based carotenoid work in cassava is an instructive template, i.e., validate HTP indices (heritability, genetic correlation, repeatability) before operational rollout. In wheat, maize, and forage grasses, modest investments in a handful of validated HTP indices delivered high per-dollar predictive performance (Lado et al., 2018; Matias et al., 2019). Operational HTP should therefore focus on low-cost, scalable platforms (ground RGB, handheld spectrometers, low-altitude UAV multispectral), automated pipelines linking images/spectra to plot/genotype IDs, and routine QC that feeds into the MT database.

Increase model complexity only as data depth supports it, and benchmark constantly. Start with additive MT-GBLUP for its interpretability and stability. As sample size and environmental diversity grow, add dominance kernels and MT×ME terms to capture non-additive variance and G×E—both influential in clonally propagated cassava and shown to improve predictions in cassava and other crops (Wolfe, Rabbi, et al., 2016; Andrade, Sousa, Oliveira, et al., 2019). Where training sets become large and heterogeneous, benchmark ML-MT methods (multi-trait deep learning, multi-kernel ensembles) against MT-GBLUP using breeder-relevant metrics (PA, RMSE, realized gain, cost). Experience from other species warns that ML yields gains only under favorable data regimes and must be carefully regularized and validated (Montesinos-López et al., 2018).

## Conclusion

This study demonstrates that MT-GP is a robust and scalable strategy for enhancing predictive accuracy, selection stability, and genetic gain in cassava breeding programs, particularly for yield-related traits. Across validation scenarios, the multivariate models (CV2–CV5) performed comparably to, or were slightly superior to, the univariate approach (CV1), maintaining predictive ability and RMSE for dry matter content while improving both metrics for fresh root yield, especially when incorporating auxiliary traits such as shoot yield and plant vigor. These traits proved particularly informative for simultaneous prediction of dry matter content and fresh root yield, reinforcing the importance of trait correlation structures in optimizing selection efficiency.

Under realistic breeding conditions, including missing-data scenarios, we confirmed the resilience and efficiency of MT models in maintaining stability and improving prediction performance across multiple traits. Although differences in classification accuracy (Kappa index) were modest, favorable auxiliary-trait combinations consistently led to higher selection responses and greater expected genetic gains, indicating tangible long-term advantages for cassava improvement. Therefore, we recommend adopting multivariate genomic prediction using auxiliary traits as an efficient and practical tool for predicting high-cost traits in cassava. Its implementation can substantially enhance the accuracy, genetic gain, and operational efficiency of cassava breeding pipelines, supporting the long-term goal of developing high-yielding, resilient, and quality-improved cassava varieties for tropical agriculture.

## Acknowledgment

The authors thank CNPq (Conselho Nacional de Desenvolvimento Cientıfico e Tecnológico), FAPESB (Fundação de Amparo à Pesquisa do Estado da Bahia), and CAPES (Coordenação de Aperfeiçoa-mento de Pessoal de Nıvel Superior) for financial support. This work was also supported by the NEXT-GEN Cassava project, through a grant to Cornell University by the Bill & Melinda Gates Foundation and the UK Department for International Development This preprint was created using the LaPreprint template (https://github.com/roaldarbol/lapreprint) by Mikkel Roald-Arbøl.

